# Top-down regulation of ingestive behavior fragmentation

**DOI:** 10.64898/2026.03.31.715709

**Authors:** Tianbo Qi, Colton Krull, Verina H. Leung, Vlad Mardare, Dong Yang, Neeraj Lal, Jingrui Ma, Shiqi Wang, Hanbing Shen, Alan Zhang, Bohan Zhao, Saba Heydari Seradj, Tatiana Korotkova, Ann Kennedy, Li Ye

**Affiliations:** Department of Neuroscience, Dorris Neuroscience Center, Scripps Research; San Diego, CA, USA; Howard Hughes Medical Institute; Chevy Chase, MD, USA; Institute for Systems Physiology, Faculty of Medicine and University Hospital Cologne, University of Cologne; Cologne, Germany; Cluster of Excellence Cellular Stress Responses in Aging-associated Diseases (CECAD), Center of Molecular Medicine Cologne (CMMC), University of Cologne; Cologne, Germany

## Abstract

In natural environments, animals rarely feed continuously to satiation; instead, feeding occurs in brief bouts separated by pauses. This fragmentation is thought to balance internal drives with external demands, yet its underlying neural mechanisms remain unclear. By combining bidirectional neural activity mapping and behavioral phenotyping, we identify a projection from the dorsal subiculum (dSub) of the hippocampus to the mammillary body (MB) as a key regulator of this fragmentation. Activity along the dSub–MB pathway tracks and gates the duration of individual feeding bouts, independent of homeostatic state. A simple bistable attractor model captures both dSub–MB neural dynamics and associated behaviors across optogenetic and behavioral perturbations. Together, these findings identify a top-down circuit mechanism that implements action selection in a naturalistic setting.

## Main

Feeding is unequivocally crucial for survival. But even when hungry, animals rarely eat continuously to satiety. Instead, feeding is often fragmented into short episodes, referred to as bouts, interspersed with pauses and other behaviors^1^. Such fragmentation of behavior, particularly common in mammals and birds, is thought to serve an important adaptive purpose, balancing feeding with expression of other behaviors necessary for predator vigilance, optimization of foraging strategies, and monitoring of social cues^2–4^. This dynamic toggling between homeostatic demands and other actions exemplifies the broader problem the brain must solve in translating internal states into moment-to-moment behaviors. Hunger and other physiological needs typically accumulate slowly and do not require an immediate reaction; they therefore shape behavior not through fast stimulus-response loops, but by preferentially biasing organisms towards some actions over others in a manner dependent on need intensity. But compared to a rich literature on how homeostatic needs are estimated, updated, and satiated^5–10^, less is known about the neural mechanisms by which need states shift this moment-to-moment organization of ingestive behavior.

Fragmented feeding is the product of an action-selection process that depends on both an animal’s internal needs and its environment. At any moment, the environment offers a range of potential action opportunities that constrain the space of behaviors an animal might produce^11,12^. Extensive experimental and theoretical work has established the hippocampus as a key site where cognitive maps of an animal’s environment, its affordances, and its reward structure are constructed^13–18^. Further work has put forth the model that a major function of hippocampus is short-term setting of goals, particularly in forecasting the consequences of a sequence of actions^19–22^. The hippocampus is therefore well poised to connect an animal’s need state and sensory environment to plans about its moment-to-moment actions.

The hippocampus is furthermore well positioned to exert top-down control over feeding behavior. Conserved across mammals and birds, the hippocampus integrates diverse external sensory features and projects to the forebrain and hypothalamus feeding network^23,24^. Rodent studies have shown that the hippocampus plays a key role in contextual feeding and is generally a negative regulator of food intake^25–31^. Moreover, human hippocampus lesion and damaged patients, including the patient H.M., have been reported to show loss of a stopping signal for eating^32,33^. We therefore explored the role of the hippocampus in the regulation of feeding microstructure.

Here we combined bidirectional activity mapping and behavioral phenotyping to identify a top-down projection from the dorsal subiculum (dSub) in the hippocampus to the mammillary body (MB) of the hypothalamus as a key circuit for regulating feeding fragmentation and bout structures. Population activity of dSub-MB projection is bimodally distributed, with “down” states closely tracking individual feeding bouts across different homeostatic states and food types. This bimodal distribution resembles a bistable system in which animals “toggle” between feeding and other behaviors depending on dSub-MB activity. Modulating dSub-MB activity specifically regulates the microstructure of feeding bouts, independent of the canonical, homeostatic control of feeding. Stochastic activations of dSub lead to transient intra-bout interruptions, and dSub-MB activity gates the transition to a full termination.

Taken together, our findings support the conclusion that the dSub projection to MB shapes moment-to-moment action selection related to feeding, rather than the underlying homeostatic drive. This raises the intriguing possibility of the subiculum as a hippocampal output that gates the execution of motivated behaviors controlled by downstream circuits, pointing to the hippocampus as a site where slow-evolving physiological needs are weighed against each other and the environment to translate them into immediate actions.

### The activity of dorsal subiculum inversely correlates with feeding bouts

The dorsal hippocampus encompasses structured areas with distinct functions and connectivity. We decided to first map the neural activity patterns across the dorsal hippocampus during natural feeding behaviors. In addition to the classic neuronal activity integrator cFos, we also incorporate an inverse activity marker pPDH, which marks the decrease of activity^34^ (Figure 1A). After overnight fasting, activities across dorsal hippocampus subfields (DG, CA3, CA1 and subiculum) generally increased, indicated either by an increase in cFos or a decrease in pPDH level (Figure 1B, 1D), consistent with a previous report^35^. One hour after re-feeding, we observed an increase of pPDH staining in the dorsal subiculum (Figure 1C) but not in other subregions, suggesting that feeding is associated with unique activity fluctuation in the dSub, motivating us to focus on this relatively underexplored area of hippocampus for its role in regulating feeding behaviors.

**Fig. 1.**
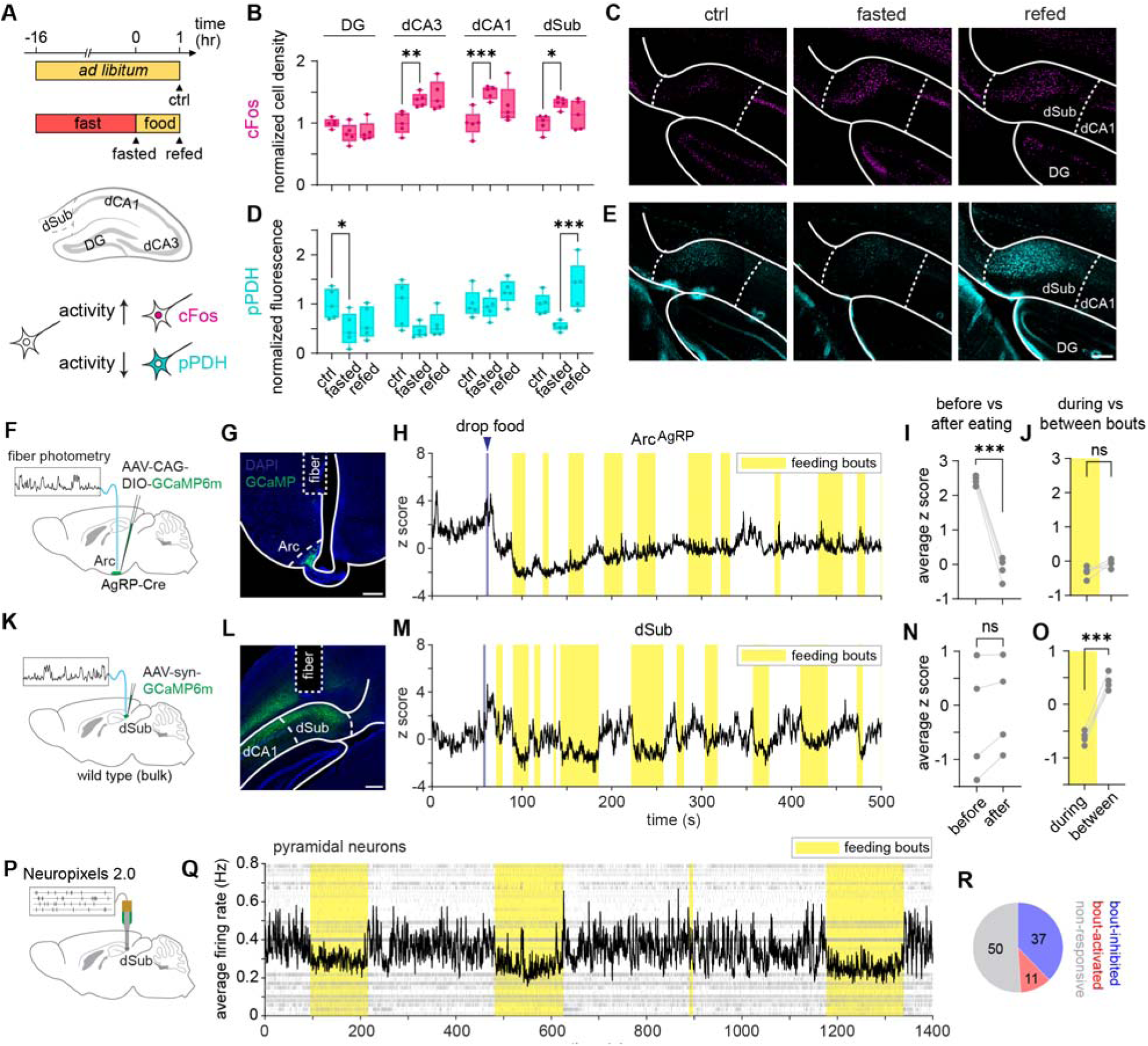
The activity of dorsal subiculum inversely correlates with feeding bouts. (A) Schematic of bidirectional activity mapping of the dorsal hippocampus in feeding behavior. (B) Quantification of cFos cell density across the dorsal hippocampus. Positively stained cells were quantified and normalized to the control group. N = 5 mice for each group. (C) Representative images of cFos staining in dSub. (D-E) Quantification of pPDH intensity across the hippocampus, and representative images of pPDH in dSub. (F-G). Schematic and representative histology of fiber photometry recording of AgRP neurons in Arc, showing virus infection and fiber placement. (H) Representative single trace of activity of Arc AgRP neurons. (I-J) Average calcium signal before (for 1 minute before food drop) vs after feeding (for the last minute of recording), and during vs between feeding bouts. N = 4 mice. (K-M) Schematic, representative histology and representative single trace of fiber photometry recording in dSub, showing virus infection and fiber placement. (N-O) Average calcium signal before vs after feeding, and during vs between feeding bouts. N = 4 mice. (P) Schematic of Neuropixels 2.0 recording in dSub. (Q) Raster plot and populational average firing rate of putative pyramidal neurons from a representative mouse. For readability given the long time window, only every 20th spike is displayed in the raster plot. (R) Proportion of bout-activated, bout-inhibited and non-responsive pyramidal neurons. Total of 98 neurons from 4 mice. Statistics determined by one-way ANOVA with Tukey’s multiple comparisons test for adjacent conditions in (B) and (D), and paired t test in (I-J) and (N-O). ∗p < 0.05; ∗∗p < 0.01; ∗∗∗p < 0.001. All scale bars are 200 μm.

Both cFos and pPDH integrate neural activity over time. To better understand the temporal relationship between dSub activity and feeding, we therefore used fiber photometry to measure calcium dynamics in dSub. To compare dSub activity with a canonical feeding circuit, we also recorded neural activity of AgRP neurons of the arcuate nucleus of the hypothalamus (Figure 1F, 1K). Unlike AgRP neurons, which show a signature decrease in activity upon the start of feeding^36^, overall dSub activity levels remained similar before and after feeding (Figure 1I, 1N). However, further examination of dSub activity revealed structured dynamics, with lower activity during feeding bouts compared to intervals between bouts (Figure 1O). This correlation of neural activity with feeding bouts was not observed in AgRP neurons (Figure 1J). This result was further confirmed by single-unit recordings using Neuropixels (Figure 1P). The population firing rate of dSub neurons decreased during feeding bouts compared to intervals between bouts. Around half of dSub neurons are modulated by feeding bouts, among which the majority show lower activity during feeding bouts (Figure 1Q-1R). Taken together, our data reveal modulation of dSub activity that correlates with the microstructure of intra-meal feeding bouts, with dynamics that are distinct from those of the classic hypothalamic circuit underlying homeostatic feeding control.

### dSub regulates feeding microstructure through its projection to the mammillary body

dSub is a major output node of the hippocampus, although its function is less understood compared to CA1^37^. To understand how dSub might contribute to the regulation of the feeding network, we mapped the efferent connections of this region. After expressing a fluorescent axon tracer (rCOMET^38^) from the dSub, we performed whole-brain tissue clearing and light-sheet microscopy to identify projections across the whole brain (Figure S3A). Fluorescence-based structural tensor analysis (STA)^39,40^ identified two main streams of extrahippocampal axonal projections: the anterior pathway that goes through the fornix and mainly terminates in the mammillary body (MB) and a group of anterior thalamic nuclei (ATN), and the posterior pathway that goes through the alveus-angular bundle and targets the entorhinal cortex (EC) (Figure S3B-S3E). Conversely, we injected the retrograde tracer cholera toxin subunit B (CTB) into the ATN, MB, and EC, and confirmed that retrograde tracing from all three regions labeled cell bodies in dSub (Figure S3G-S3H). Hence, our anterograde and retrograde anatomical tracing revealed the major extrahippocampal projections from dSub, largely consistently with the literature^41,42^.

To test whether any of these dSub projections regulate feeding behavior, we performed a projection-specific terminal photoinhibition using the light-activated inhibitory GPCR eOPN3^43^. We expressed eOPN3 in dSub and implanted an optic fiber above MB, ATN, or EC to selectively inhibit each dSub projection pathway (Figure 2A-2B, S4C, S4G, S4K). None of the dSub projection inhibitions changed the overall amount of food intake (Figure 2D, 2G, 2J), whereas similar eOPN3-based photoinhibition in a well-established feeding center, the lateral hypothalamus (LH), resulted in an apparent decrease (Figure S4A-S4B, used as a technical control^44–46)^. But interestingly, when we quantified the statistical properties of the feeding bout microstructure, we found that inhibition of the dSub-MB pathway, but not the other two, increased the duration of feeding bouts, as indicated by an increase in mean bout duration and a right shift of the distribution of bout duration (Figure 2E, 2H, 2K). Inter-bout intervals and latencies remained unchanged in all three cohorts (Figure 2F, 2I, 2L, S4C-S4N), suggesting that the dSub-MB pathway affects bout terminations, but not initiations.

**Fig. 2.**
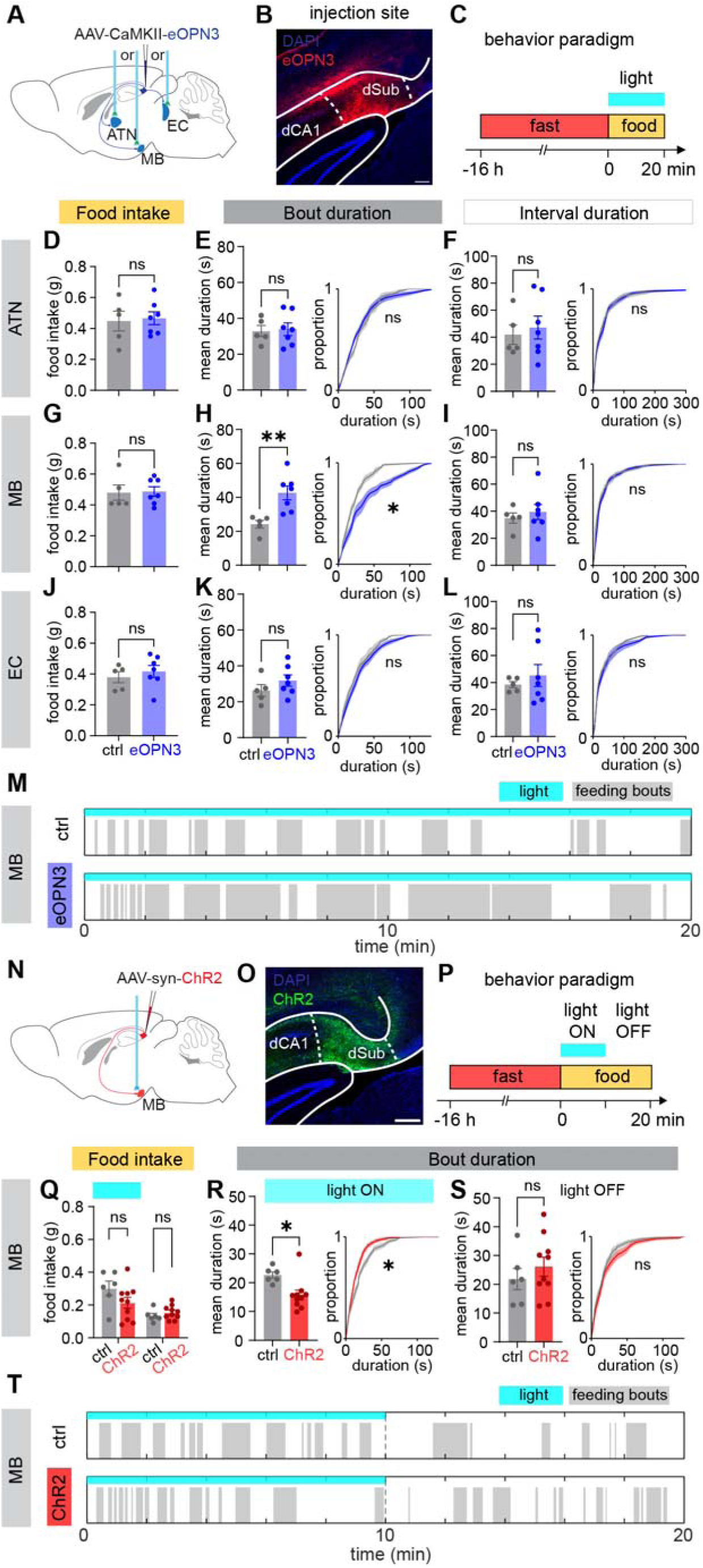
dSub regulates feeding microstructure through its projection to the mammillary body. (A) Schematic of virus injection and fiber implant for photoinhibition experiments. (B) Representative histology showing eOPN3 expression in dSub. (C) Schematic of the fast-refeeding photoinhibition experiment. (D) Food intake analysis under dSub-ATN photoinhibition. (E-F) Bout duration and interval duration analysis under dSub-ATN photoinhibition. Left: mean duration; right: empirical cumulative distribution function (ECDF) of duration. N = 5 mice for ctrl group and N = 7 mice for eOPN3 group. (G-I) Food intake, bout duration and interval duration analysis under dSub-MB photoinhibition. N = 5 mice for ctrl group and N = 7 mice for eOPN3 group. (J-L) Food intake, bout duration and interval duration analysis under dSub-EC photoinhibition. N = 5 mice for ctrl group and N = 7 mice for eOPN3 group. (M) Representative raster of feeding bouts of a ctrl mouse and an eOPN3 mouse during dSub-MB photoinhibition. (N-O) Schematic of virus injection and fiber implantation for photoactivation experiments, and representative histology showing ChR2 expression in dSub. (P) Schematics of the fast-refeeding photoactivation experiment. (Q) Food intake in light ON phase and light OFF phase of dSub-MB photoactivation. (R-S) Bout duration analyses in light ON phase and light OFF phase of dSub-MB photoactivation. N = 6 mice for ctrl group and N = 10 mice for ChR2 group. (T) Representative raster of feeding bouts of a ctrl mouse and a ChR2 mouse during dSub-MB photoactivation. All values are mean ± SEM. Statistics determined by Student’s t-test for mean duration and food intake analyses, two-tailed permutation Kolmogorov–Smirnov test (KS test) for ECDF analyses for photoinhibition experiments. Afterwards, one-tailed permutation KS test was used to test the gain-of-function hypothesis for photoactivation experiments. ∗p < 0.05. All scale bars are 200 μm.

We next optogenetically activated the dSub-MB projection using ChR2 (Figure 2N-2O). As with photoinhibition, food intake was not altered (Figure 2Q; using AgRP as a positive control (Figure S5A-S5B)). However, the effect on feeding fragmentation mirrored that of photoinhibition: feeding bouts were significantly shorter during the photoactivation session, while inter-bout intervals and latencies remained unchanged (Figure 2R, S5D-S5H). No bout duration or interval difference was found in the unstimulated light off session (Figure 2S, S5D-S5H). In addition, no general preference or aversion was associated with dSub-MB activation (Figure S5I). Together, these results indicate that the dSub-MB activity bi-directionally regulates the structure of feeding bouts without affecting the amount of food intake.

### dSub-MB activity specifically tracks individual ingestion bouts

After establishing the causal significance, we further measured the endogenous neural activity of the dSub-MB projection using fiber photometry (Figure 3A-3B). Similar to bulk dSub activity (Figure 1M), MB-projecting dSub neurons showed a time-locked decrease of activity at the initiation of each feeding bout, and an increase of activity at the termination (Figure 3C-3E), consistent with a role in feeding bout modulation. This activity pattern was consistent in both males and females (Figure S6A-S6B).

**Fig. 3.**
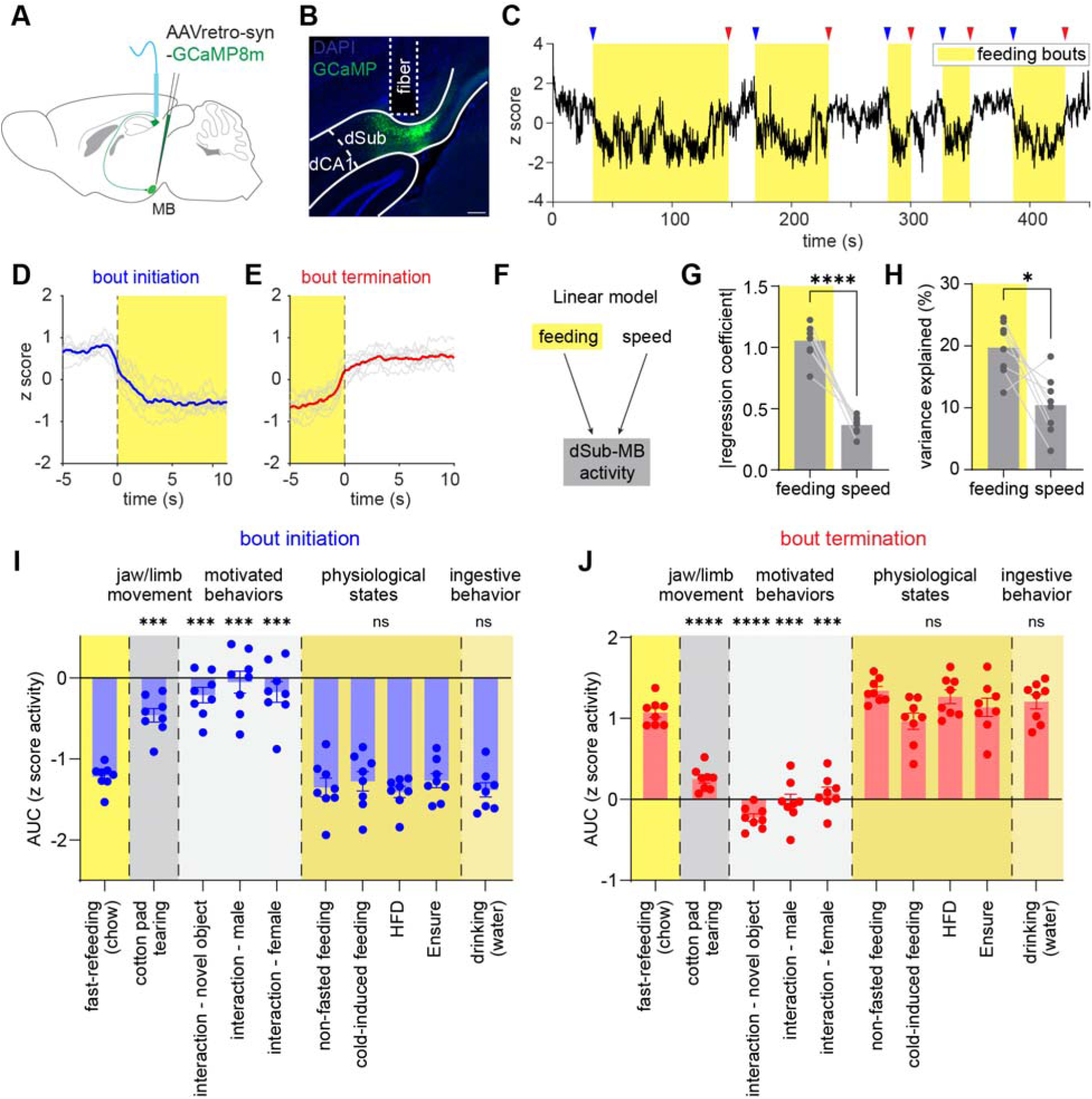
dSub-MB activity selectively tracks individual ingestion bouts. (A-B) Schematic of virus injection, fiber implantation for dSub-MB fiber photometry experiment, and representative histology showing virus expression and fiber placement. (C) Representative calcium trace of dSub-MB activity along with feeding bouts. Blue and red arrows represent the initiation and termination of feeding bouts. (D-E) Averaged calcium response aligned on initiation and termination of feeding bouts. Gray traces show the averaged response of individual mice, and blue or red traces show the averaged across mice. N = 8 mice. (F) Schematic of the linear regression of dSub-MB calcium activity against feeding bouts and speed of locomotion. (G-H) Absolute value of regression coefficient and percentage of variance explained of dSub-MB activity by feeding and speed. (I-J) AUC analysis of calcium signal at initiation and termination of bouts of different behaviors. Values are mean ± SEM in (I-J). Statistics determined by paired t test in (G-H), and repeated measures one-way ANOVA with Dunnett’s multiple comparisons test in (I-J), all compared against fast-refeeding (chow) group. ∗p < 0.05, ∗∗∗p < 0.001, ∗∗∗∗p < 0.0001. Scale bar in (B) is 200 μm.

Because hippocampal neuronal activity is known to correlate with locomotor speed^47,48^, we tested whether the correlation between dSub-MB activity and feeding was confounded by locomotion. A multiple linear regression model showed that feeding bouts exhibited a stronger correlation with dSub-MB activity and accounted for a greater proportion of variance (Figure 3F-3H), indicating that feeding bouts are the predominant driver of the observed dSub-MB activity pattern.

We next tested whether the observed neural activity could be confounded with other forms of movement, such as jaw or forelimb activity. To this end, we recorded dSub-MB activity while mice tore apart a cotton pad to make a nest, a behavior that involves a similar motor repertoire of jaw and forelimb movements to feeding. In contrast to feeding bouts, cotton pad-tearing was not associated with changes in dSub-MB activity, and neural activity was even less correlated with bouts of pad-tearing than with locomotion speed (Figure 3I-3J, S6C-S6E). These results further demonstrate that dSub-MB activity tracks ingestion bouts instead of general motor behaviors. In addition, other motivated behaviors, such as investigation of a conspecific or a novel object, were not accompanied by changes in dSub-MB activity (Figure 3I-3J). Taken together, these results strongly indicate that dSub-MB activity is selectively correlated with ingestive bouts.

Next, we tested whether dSub-MB activity is modulated by metabolic state and food type. Neither fasting nor cold-induced energy deficit^49^ altered the dSub-MB activity patterns associated with feeding bouts (Figure 3I-3J). Furthermore, during fast-induced feeding, the amplitude of calcium dynamics from early to late bouts did not change as the meal progressed (Figure S6F-S6G), suggesting that dSub-MB activity tracks feeding bout engagement rather than the hunger versus satiety state. In addition, feeding mice with different types of food, including a 60% high-fat diet and a liquid food (Ensure), resulted in similar dynamics of dSub-MB compared to a regular chow (Figure 3I-3J), indicating that the signal was not determined by caloric content, palatability, or physical form of the food. Indeed, we found water drinking was also accompanied by a similar pattern of dSub-MB activity (Figure 3I-3J). Together, these findings demonstrate that dSub-MB dynamics faithfully track individual ingestive bouts, independent of the animals’ metabolic state or the ingested content.

### dSub-MB gates transition from feeding interruption to bout termination

During feeding bouts, we sometimes noticed brief calcium transients that lasted for a few seconds (Figure 4A). Upon video examination, we found that these small events were associated with brief interruptions in feeding. To slow down feeding behavior for detailed examination, we adopted a paradigm in which mice were provided with a piece of pasta^50,51^. We found that calcium transients were associated behaviors such as brief looking around or moving before immediately resuming eating (Movie S1). We annotated these “intra-bout interruptions” in our full dataset and found that they were consistently associated with short calcium events, in contrast to the persistent rise in activity associated with bout termination (Figure 4B-4C). The distribution of time intervals between interruptions was roughly log-normal (Figure S7B). Interestingly, time intervals between bout terminations and the preceding interruption followed the same distribution (Figure S7C-S7D). This is consistent with feeding interruptions and terminations both arising from a common underlying renewal process, with each event in the process eliciting either a brief pause or a full termination of feeding.

**Fig. 4.**
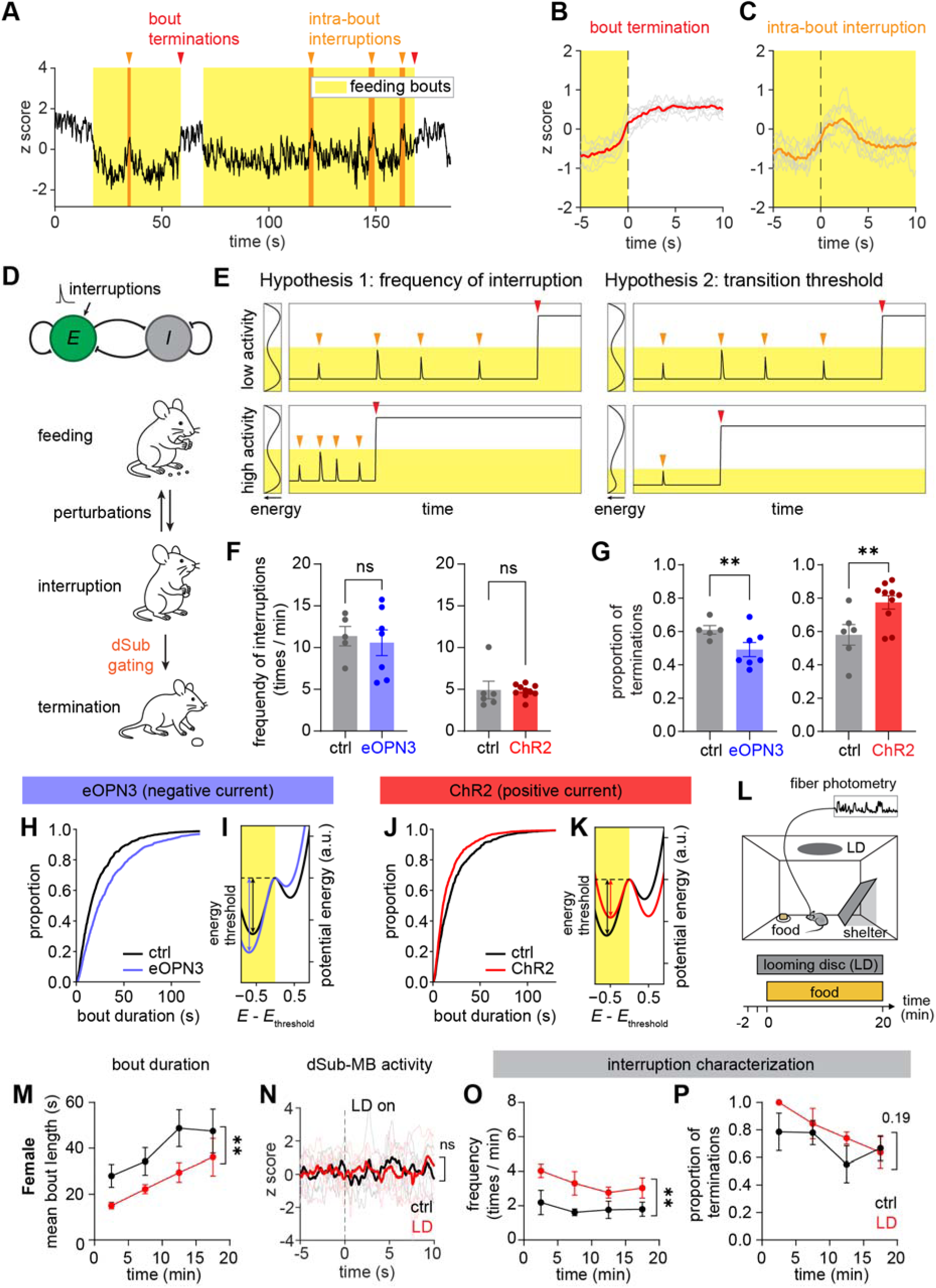
dSub-MB promotes transition from intra-bout interruption to termination. (A) Representative trace of dSub-MB activity showing bouts and intra-bout interruptions. (B-C) Averaged calcium activity aligned on bout terminations and intra-bout interruptions. N = 8 mice. (D) Top: Model of dSub as a recurrently connected network of an excitatory population (*E*) and an inhibitory population (*I*). Interruptions and optogenetic currents were modeled as inputs to the excitatory population. Bottom: Schematic of hypothesized transition structure of behavior during feeding. Interruptions are driven by a random renewal process, and dSub-MB state determines whether the interruption will fully terminate the ongoing bout. (E) Two hypotheses of how statistics of interruption inputs to dSub-MB neurons change bout length. Hypothesis 1: shorter bouts reflect higher frequency of interruption inputs. Hypothesis 2: shorter bouts reflect lower threshold to transition to non-feeding state. (F) Total frequency of interruptions and terminations in the eOPN3 cohort and the ChR2 cohort. eOPN3 cohort: N = 5 mice for ctrl group and N = 7 mice for eOPN3 group; ChR2 cohort: N = 6 mice for ctrl group and N = 10 mice for ChR2 group. (G) Proportion of terminations among all interruption and termination events. Same N as in (F). (H) Analysis of model-generated feeding bouts, showing that a tonic inhibitory input to the *E* population increases bout duration. (I) Potential energy surface of the approximated one-dimensional effective system (see Methods) in the presence (blue) or absence (black) of negative current injection to *E* population. Arrows represent the threshold energy to transition from the feeding state to the non-feeding state. (J) Analysis of model-generated feeding bouts, showing that a tonic excitatory input to the E population increases bout duration. (K) Potential energy surface of the one-dimensional effective system in the presence (red) or absence (black) positive current injection to *E* population. (L) Schematic of the looming disc experiment. (M) Time course of the bout duration in female mice. N = 5 mice for ctrl group and N = 5 mice for LD group. (N) Calcium activity of dSub-MB in female mice triggered on the first appearance of the looming disk. (O-P) Time course of the total frequency of interruptions and terminations, and percentage of terminations, in female mice. Values are mean ± SEM for (F-G), (M) and (O-P). Statistics determined by Student’s t-test for (F), and mixed effect Bernoulli generalized linear model (GLM) for (G), and two-way ANOVA (main effect of group) for (M-P). ∗p < 0.05; ∗∗p < 0.01. P values for non-significant comparisons are labeled in (M-P).

Our imaging of the dSub-MB projection is therefore suggestive of a bistable system, with a low-activity attractor during feeding and a high-activity attractor during other behaviors. Stochastic perturbations to this system during feeding can either resettle into the feeding attractor (interruptions) or transition the system into the non-feeding attractor (terminations). To formalize this intuition, we built a simple mathematical model and showed that it could recapitulate much of the phenomena associated with dSub-MB activity. Bistability is widespread in nonlinear dynamical systems, and can arise in neural networks via diverse cell-intrinsic, single population, and multi-population mechanisms. For our model we selected the Wilson-Cowan model^52^ of interacting excitatory (*E*) and inhibitory (*I*) neuron populations, a well-studied system with bistable dynamics, though we note that similar behavior can be produced via many other means, and do not claim the *E*-*I* circuit should be taken literally as a model of dSub circuit architecture.

We constructed a Wilson-Cowan model of a single pair of *E* and *I* populations and assigned low and high activity of the model’s *E* population to represent feeding and non-feeding states, respectively (Figure 4D). We modeled intra-bout interruptions as a renewal process, where each interruption consists of a pulse of input to the model *E* population, with amplitude sampled from a Gaussian distribution with nonzero mean. When the system is in the low-activity feeding state, input pulses produce transient increases in the population activity. Sufficiently large inputs will push the system out of the feeding attractor, thus ending a feeding bout (Figure S8B). Both the *E* and the *I* populations show modulation by feeding bouts, consistent with our Neuropixels recording (Figure S8C).

Tuning the event rate and amplitude of the renewal process input allowed us to tune our model such that it produced feeding bouts with statistical properties matching the real behavior data (Figure S8H, S8J). Consideration of the model then suggested two hypotheses for how perturbations to the dSub-MB projection could change feeding bout structure: neural activation might alter (1) the rate of interruption pulses, or (2) their amplitude relative to the energy threshold between states (Figure 4E).

To test these hypotheses, we re-analyzed our optogenetic manipulation data of both eOPN3 and ChR2. We found that neither optogenetic activation nor inhibition affected the total frequency of interruptions and terminations (Figure 4F), and that the renewal statistics of interruption and termination bouts were unchanged by either manipulation (Figure S8D-S8G), disputing the first hypothesis. But strikingly, the ratio of terminations to interruptions increased during ChR2-mediated activation and decreased during eOPN3-medinated inhibition (Figure 4G). These results support the second hypothesis, that stimulation of dSub-MB does not alter the rate of interruptions; rather, it increases the likelihood that interruptions will transition to full termination of a feeding bout.

Finally, we showed that a simple way to match these effects in the model is by current injection into the model neural populations. Specifically, injection of negative (inhibitory) current into the *E* population increased the energy threshold and shifted the model towards longer bouts in the feeding state, matching the experimental result of the eOPN3-mediated inhibition (Figure 4H-I). By contrast, injection of positive current to the *E* population reduced the energy threshold and shifted the model to shorter bout durations, matching the ChR2-mediated activation results (Figure 4J-K). Thus our simple model can match both the neural activity, the animal behavior, and the result of circuit manipulation in the dSub-MB projection. Altogether, our experimental data and mathematical model provide a robust framework to understand how control of feeding fragmentation can be achieved by tuning the energy threshold between states in a bistable system.

### Visual threat shortens feeding bouts by increasing interruptions

Our model succinctly explains how photoactivation and inhibition of dSub-MB alter feeding structures by changing the energy threshold between attractor states of a neural network. Intriguingly, the model also predicts that it should be possible to shorten feeding bouts by increasing the interruption rate without changing the gating threshold. The model further predicts that a perturbation that changes the rate of interruptions should leave the level of dSub-MB neural activity in the feeding and non-feeding states unchanged. To test whether this prediction holds in real experiments, we sought alternative means of inducing feeding bout fragmentation. We found that an external threat cue—the well-established looming disc paradigm^53–55^—was capable of perturbing feeding behavior by increasing animals’ vigilance (Figure 4L). Presentation of a looming disc prior to food introduction led to shorter feeding bouts, although the reduction was more prominent and longer lasting in females compared to male mice (Figure 4M, S9A). Quantification of latency to the first feeding bout confirmed the presence of sex differences, again with a more pronounced phenotype in females (Figure S9E-S9F).

Interestingly, the looming disk exposure did not directly alter dSub-MB activity in either sex (Figure 4N, S9B). Thus, consistent with our model’s prediction, the shortening of feeding bouts in the presence of the looming disc was characterized by an increase in the total frequency of interruptions (Figure 4O, S9C) without altering the proportion of terminations (Figure 4P, S9D). Altogether, these findings demonstrate that our model can explain the top-down regulation of feeding fragmentation in both experimental and naturalistic settings.

## Discussion

In this study, we identified a top-down hippocampal-hypothalamic circuit that supports the natural fragmentation of feeding behaviors. Within each feeding bout, transient input to the dSub-MB pathway perturbs the neural state, and dSub-MB activity gates the likelihood of behavioral transitions. This circuit mechanism is distinct from the canonical homeostatic regulation of feeding, as it operates independently of metabolic state and caloric intake. More broadly, our results indicate that hippocampal output directly regulates the stochasticity of moment-to-moment action switching, thereby enabling flexible transitions between competing actions in response to internal needs and environmental contingencies. This finding echoes the recent findings that the hippocampus underlies trajectory planning in navigation tasks^56,57^, and extends its functional role to general action selection beyond spatial navigation.

In addition to its well-known role in memory and spatial navigation, the hippocampus has been implicated in feeding control. Earlier studies in humans and rodents showed that the hippocampus negatively regulates food intake^25,26,32,33,58–60^. Recent works have dissected several hippocampal outputs, including projections to the septum, the lateral hypothalamus, and the nucleus accumbens, that regulate feeding, mainly in the paradigm of contextual feeding^27–30,61,62^. Our work complements these findings to support a unifying role for the hippocampus in integrating multisensory information to exert top-down control over broader behaviors. Within this view, the dSub-MB circuit serves as a major hippocampal output that translates an integrated hippocampal cognitive map into downstream behavioral control. More broadly, this regulation of ingestion bout termination might represent a general role of the hippocampus in the transition from consummatory behaviors (hippocampal non-theta states-associated) to preparatory or appetitive behaviors (theta states-associated)^63,64^, although the direct test of this hypothesis is beyond the scope of the current study. Furthermore, the exact source of perturbation input to the circuit, which likely arises from the upstream hippocampal-entorhinal network, remains to be further investigated.

The connection between the hippocampus and MB was described as early as in 1937 as a component of the “Papez circuit”^65^, originally proposed to mediate emotional processing but still poorly understood^66^. Previous studies have shown that MB is anatomically connected to the reticular formation in the brainstem^67–69^, where reside neurons that mediates arousal and motor functions. These connections suggest that the dSub-MB pathway may promote a heightened arousal state, allowing animals to interrupt feeding bouts and remain responsive to external stimuli.

Setting priorities between competing physiological and environmental cues is critical to survival: animals must eat to meet their nutritional needs, but not at the cost of missing signs of an approaching predator. It is rarely the case that a single external cue or physiological drive will fully command an animal’s behavior, driving urgent responses to an imminent survival threat. Instead, most of the time, animals face a variety of needs of varying urgencies, and they structure their actions to balance all of these needs in parallel, freely toggling between self-tending, ingestive, social, and vigilance behaviors. The top-down regulation of behavioral fragmentation we uncover here reveals a core feature of feeding, namely its organization into bouts, that has been largely overlooked by the feeding literature because it reflects not the intensity of an animal’s drive to eat but how the animal balances this with other tasks of living. Thus our work provides a key insight into how the drive to eat informs the broader problem of action selection in naturalistic settings.

## Acknowledgments

We thank C. Ran and E. Zorilla for behavior setups. We thank all members of the Ye lab and the Dorris Neuroscience Center for their support and feedback.

## Author contributions

Conceptualization: LY, TQ; Methodology: LY, AK, TQ; Investigation: TQ, CK, VHL, VM, DY, NL, JM, SW, HS, AZ, BZ, SHS; Funding acquisition: LY; Project administration: LY, AK, TK; Supervision: LY; Writing – original draft: LY, AK, TQ.

## Data, code, and materials availability

All data necessary to understand the conclusions of this study are available in the main text and supplementary materials. Code for fluorescence-based STA is available on GitHub (https://github.com/The-Ye-Lab/FluoSTA). Code for modeling of dSub-MB circuit is available on GitHub (https://github.com/The-Ye-Lab/dSub-manuscript).

## Materials and Methods

### Animals

All animal experiments were performed in accordance with the National Institutes of Health Guide for the Care and Use of Laboratory Animals and approved by Scripps Research. C57BL/6J mice (Jackson Laboratory; C57BL/6J; stock #000664) and AgRP-Cre mice (Jackson Laboratory; Agrptm1(cre)Lowl/J; stock #012899) were used. For wild-type mice, male mice were used for activity marker staining and optogenetics experiments, and both sexes were used for fiber photometry experiments; for AgRP-Cre mice, both sexes were used. Mice were group housed and maintained on a 12 h light: 12 h dark cycle with food and water *ad libitum* unless specified. Mice were assigned to experimental groups based on balanced body weight.

### Viral injections and stereotaxic surgery

AAV9-Syn-GCaMP6m-WPRE-SV40 (Addgene #100841, 7E+12 vg/mL), AAV9-CAG-Flex-GCaMP6m-WPRE-SV40 (Addgene #100839, 1.3E+13 vg/mL), AAV1-CaMKIIa(0.4)-eOPN3-mScarlet-WPRE (Addgene #12712, 3E+12 vg/mL), AAV1-CaMKIIa-mCherry (Addgene #114469, 3E+12 vg/mL), AAV9-hSyn-hChR2(H134R)-eYFP (Addgene #26973, 3E+12 vg/mL), AAV9-hSyn1-eYFP (Addgene #117382, 3E+12 vg/mL), AAV9-EF1a-DIO-hChR2(H134R)-EYFP-WPRE-HGHpA (Addgene #20298, 2.3E+13 vg/mL), AAVretro-syn-jGCaMP8m-WPRE (Addgene #162375, 1.9E+13 vg/mL) were obtained from Addgene. AAV8-CaMKIIa-rCOMET (3.6E+13 vg/mL) was obtained from the Stanford viral core. The viruses were diluted to the respective titer using DPBS. CTB-488, CTB-594 and CTB-647 (C22841, C34777, C34778, Invitrogen) were diluted to 0.2% (w/w) using DPBS.

Mice were anaesthetized with 1.5% isoflurane. The skull was mounted onto a stereotaxic frame (David Kopf Instruments) and balanced using the bregma and lambda system. The virus was infused at 100 nL min⁻¹ through a glass pipette with a nanoinjector (Nanoliter 2020 Injector, 300704, World Precision Instrument). The pipette was kept at the injection site for 5-10 min before withdrawing. Mice were allowed to recover for at least 10 days before experiments. For fiber implant, fiber optics (200 μm core, 0.39 NA for optogenetics, and 400 μm core, 0.50 NA for fiber photometry) were lowered to the site, and the cannulae were then fixed onto the skull using dental cement (C&B-Metabond, Parkell).

The following coordinates were used for injections: dSub (A/P −3.7 mm, M/L 2.15 mm, D/V 1.8 mm), Arc (A/P −1.6 mm, M/L 0.3 mm, D/V 5.7 mm), MB (A/P −2.7 mm, M/L 0.15 mm, D/V 5.2 mm), LH (A/P −1.5 mm, M/L 1.0 mm, D/V 5.0 mm). The following coordinates were used for fiber implants: dSub (A/P −3.7 mm, M/L 2.15 mm, D/V 1.6 mm), Arc (A/P −1.6 mm, M/L 0.2 mm, D/V 5.5 mm), ATN (A/P 1.0 mm, M/L, 1.25 mm, 2.0 mm, with a 10° angle), MB (A/P 2.7 mm, M/L 0.0 mm, D/V 5.0 mm), EC (A/P −4.7 mm, M/L 3.5 mm, D/V 2.9 mm), LH (A/P −1.5 mm, M/L 1.0 mm, D/V 4.8 mm, with a 10° angle). All coordinates were relative to the Bregma point. The following injection volume were used for each experiment: 250 nL in dSub for dSub fiber photometry, 500 nL in Arc for AgRP fiber photometry, 300 nL in dSub for anterograde tracing from dSub, 500 nL in ATN, MB and EC for CTB retrograde tracing to dSub, 150 nL in dSub for dSub projection-specific optogenetics experiments, 300 nL in LH for LH optogenetics experiment, 500 nL in Arc for AgRP optogenetics experiment, 500 nL in MB for dSub-MB fiber photometry.

### Behaviors

For fasting–refeeding experiments, mice were food-deprived for 16 h prior to testing. During recording, mice were placed in a tall chamber, and the implanted cannula was connected to the fiber photometry setup. After a 10 min habituation period, a piece of food (standard chow or HFD) or a water bottle containing Ensure was introduced, and calcium signals together with behavior were recorded for 30 min. In the non-fasted feeding condition, mice had *ad libitum* access to food prior to the session and were recorded for 1 h with a piece of chow. For thirst–drinking experiments, mice were water-deprived for 16 h, provided access to water, and recorded for 10 min. Cold-induced feeding was conducted as previously described ^49^, and nesting behavior under cold conditions was examined similarly, with a cotton pad provided. For investigation assays, a novel object (15 mL Eppendorf tube cap), a male mouse, or a female mouse was placed in the chamber, and behavior was recorded for 10 min.

Video recording of behavior was used to manually annotate and timestamp feeding and other behaviors using a behavior annotator ^70^. Intra-bout interruptions were defined as feeding intervals shorter than 4 s (see Figure S7A).

For speed analysis, the position of the mouse was tracked with DeepLabCut ^71^ for the calculation of speed for regression analysis.

### Fiber photometry

Mice were attached to a patch fiber (400 μm core, 0.57 NA, Doric lenses) connected to 470 and 410 nm light sources. Fiber photometry acquisition setup and the pre-processing have previously been described ^72^. The reference-subtracted signal was z-scored and aligned to the behavior events. For AUC analysis, the average signal within 10 s before the event was used as the baseline, and the average signal of min{20 s, behavior duration} after the event was calculated as AUC.

### Electrophysiology

Mice were anesthetized with isoflurane and placed in a stereotaxic frame. The skull was cleaned, dried, and marked with stereotaxic coordinates. A Neuropixels probe ^73^ was assembled using a custom-made holder ^74^. The implant was slowly lowered into the dorsal subiculum (A/P −3.08 mm, M/L 1.50 mm, D/V 1.60 mm) at a rate of 2 µm s⁻¹. Fast-refeeding recording experiments were performed similar to fiber photometry recording experiments. Recordings were performed using a setup enabling the use of Open Ephys and synchronized with the DaqBox^75^. Signals were acquired continuously at 30 kHz using Open Ephys ^76^ and synchronized with video recordings of behavior. The combined files were processed with Kilosort4 ^77^ and subsequently manually curated in Phy2 ^78^ based on areas derived from the anatomical reconstruction of the probe position. Relevant outputs were extracted through a custom Python script and neurons were manually classified into pyramidal or interneuron based on spike waveform shape ^42,79,80^. TTP was defined as the time between the minimum of the second peak and the maximum of the third peak within the action potential waveform with the threshold being 0.4. These neurons were further classified as inhibited or excited based on their activity during feeding bouts: neurons whose average firing rate during feeding bouts were significantly higher (lower) than bout intervals were considered as bout-activated (-inhibited) neurons.

### Optogenetics

Mice were connected to a light source with an optical patch cord (200 μm core, 0.37 NA, Doric lenses). For eOPN3 inhibition experiments, green LED light (554 nm, MINTF4, Thorlabs) at 2.5 mW power was delivered in 10 ms pulses at 50 Hz ^81^. Mice were then acclimated to the behavior chamber for 10 min, and then provided with a piece of chow, and recorded for 20 min with photostimulation. For ChR2 activation experiments, blue laser light (473 nm, Intelligent Optogenetics System, RWD) at 12 mW power was delivered in 10 ms pulses at 20 Hz. Mice were recorded for 10 min with photostimulation, and then 10 min without. Food intake was manually measured by the food weight before and after each session.

For the RTPP experiment, mice were placed in a two-chamber acrylic box (60 × 25 × 30 cm), each side having different pattern on the wall. The location of the mouse was tracked and used to control the optogenetic machine using the Track-Control toolbox ^82^. One side was paired with optogenetic stimulation, and this side was randomized across mice. A total of 30 min was recorded and analyzed.

### Looming disc experiment

Mice expressing GCaMP and implanted with an optical fiber were connected to the fiber photometry recording system and placed in a tall chamber. A chow pellet was fixed in one corner of the chamber and covered with a metal mesh to prevent access. A shelter was positioned in another corner. Mice were allowed to acclimate to the chamber for 1 min prior to the start of recording. For the looming disc (LD) group, continuous looming disc stimuli ^53^ (black disc on a gray background; ∼20° visual angle; 250 ms expansion, 250 ms hold at maximum size, 500 ms interval) were presented from an overhead screen beginning 1 min after recording onset. 1 min after the onset of the looming stimulation, the metal mesh covering the food was removed, allowing access to the chow. Mice were then allowed to freely consume the food for 20 minutes. Behavioral responses were video recorded and subsequently analyzed.

### Histology and immunohistochemistry

Mice were terminally anesthetized with isoflurane and intracardially perfused with PBS and 4% PFA. Brains were dissected and post-fixed in 4% PFA overnight at 4 °C. Brains were then dehydrated in 30% sucrose for 24 h and embedded in OCT (Tissue-Tek O.C.T. Compound, Sakura). Coronal sections were prepared using a Cryostat (FS800A, RWD). 80 µm sections were cut for histology without staining, and 50 µm sections were cut for immunostaining.

For immunostaining of cFos and pPDH, free-floating sections were blocked with blocking buffer (5% normal donkey serum in PBS with 0.3% Triton X-100 (PBST)) at room temperature for 1 h, and then incubated with primary antibodies (rabbit monoclonal anti-pPDH (Ser293) (Cell Signaling #37115), mouse monoclonal anti-c-Fos (Santa Cruz Biotechnology #sc-271243), both 1:500 dilution) in blocking buffer at 4 °C overnight. Slices were then washed by PBST for 3 × 15 min, and incubated with secondary antibodies (Donkey Anti-Mouse IgG Antibody (Alexa Fluor 488), Donkey Anti-Rabbit IgG Antibody (Alexa Fluor 647), Jackson ImmunoResearch, both 1:500 dilution) in blocking buffer at room temperature for 1 h. After washing with PBST for 3 × 15 min, slices were stained with DAPI (5 nM in PBS) for 30 min, and then mounted in fluoromount-G (Electron Microscopy Science) for imaging using a slide scanner (VS200, Olympus) or a confocal microscope (FV3000 or FV4000, Olympus).

For quantification of cFos, positive stained cells were identified with a customized ImageJ macro script using the thresholding and particle analysis function. For pPDH intensity, the average intensity of the whole region was quantified. The cell density and intensity were averaged, weighed by area, across multiple sections of the same mouse to obtain the final value.

### Tissue clearing, lightsheet imaging, and STA

Whole brains were cleared using the HYBRiD protocol ^83^. Cleared whole brains were RI-matched in EasyIndex (RI = 1.52, LifeCanvas Technologies) overnight, mounted in 1% agarose in EasyIndex for imaging, and then equilibrated overnight in the immersion oil (RI = 1.52). Imaging was performed using a lightsheet microscope (SmartSPIM, LifeCanvas Technologies) with a 3.6x objective, 0.28 NA, with imaging resolution 1.8 µm, 1.8 µm, 2 µm, xyz voxel size. Raw images were then destriped and stitched for further processing. Images were visualized in IMARIS.

The images were then downsampled by 2x, and median-filtered on the xy plane to remove the stripes on the z axis. Fluorescence-based STA were performed using customized scripts adapted from MIRACL ^39^. Seeds were manually drawn in IMARIS. In brief, the third eigenvector field of the Hessian matrix was used to track the streamlines from the seed. Streamlines were filtered by turning angle, length, image intensity and coherence. Streamlines were visualized in MRview.

### Computational modeling

The dSub-MB circuit was modeled using a stochastic Wilson-Cowan model ^52^ with an excitatory population and an inhibitory population, following

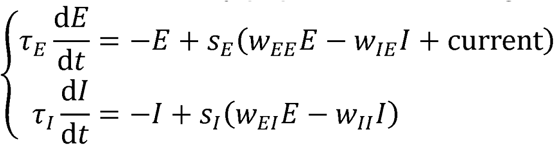

Where τ_E_ = 2s and τ_I_ = 0.4s are the time constants, and w_EE_ = w_II_15 and w_IE =WEI = 10_ are the synaptic strengths. Current was set to −0.7 and 0.85 for eOPN3 and ChR2, respectively.s_E_ and s_I_ 626 are the gain functions of the populations, following

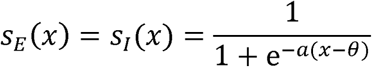

where a = 1 and θ 8. At each interruption, E was updated according to E ← E + Δ where the pulse Δ ∼ N (0.560.11^2^) is the transient input. Interruptions follow a renewal process whose interval follows a lognormal distribution fitted from data, log(interval/s) ∼ N(2.22,0.99^2^) Gaussian noise with a mean of 0 and a standard deviation of 0.1055 was added to the equations to capture stochastic variability. The equations were simulated using the Euler–Maruyama method with a time step of 0.02 s.

For quasi-static approximation, because τ_I ≪_ τ_E_ I quickly converges to its steady state at given *E*. The steady state *I*, denoted Î (E), was calculated through solving dI / dt = 0. This Î (E) was then plugged into the dE/dt equation to obtain a 1-dimensional system of E

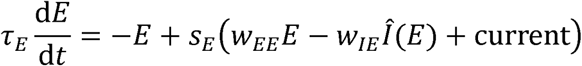

Then, the potential energy V is calculated as the negative integral of dE /dt, zeroed at the threshold of E

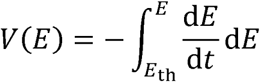

### Statistics

Statistical analysis was performed using Prism 10 (GraphPad) for t tests and ANOVA, Python scripts for permutation KS tests, and R or Python scripts for GLMs. All statistical methods and numbers of biological replicates are indicated in the corresponding figure legends. P value of 0.05 or less was considered statistically significant.

---

**Fig. S1.**
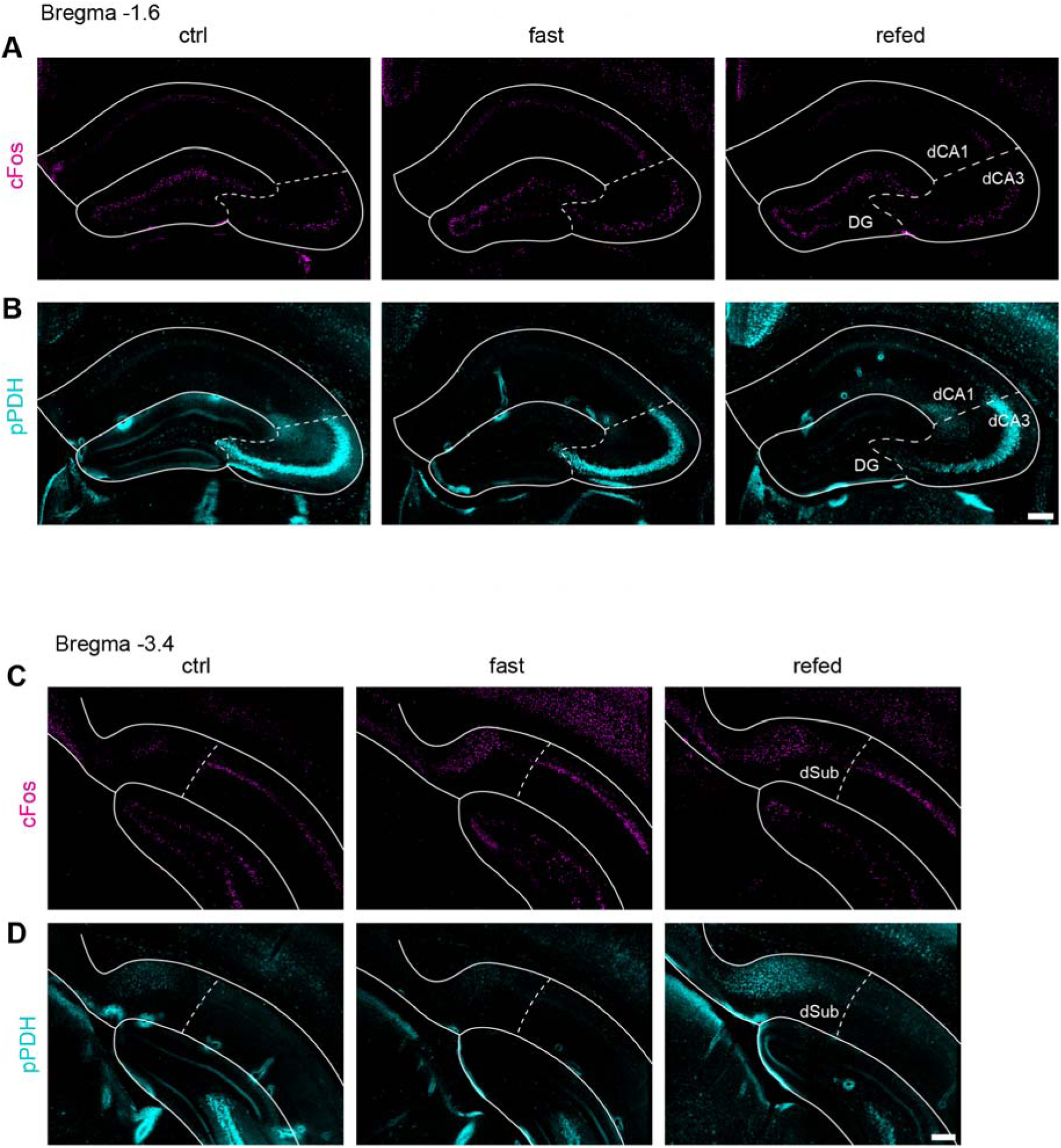
Bidirectional activity mapping of the dorsal hippocampus. (A) Representative images of cFos staining in DG, CA1 and CA3. (B) Representative images of pPDH staining in DG, CA1 and CA3. (C) Representative images of cFos staining in posterior part of the dorsal hippocampus, showing dSub. (D) Representative images of pPDH staining in posterior part of the dorsal hippocampus, showing dSub.

**Fig. S2.**
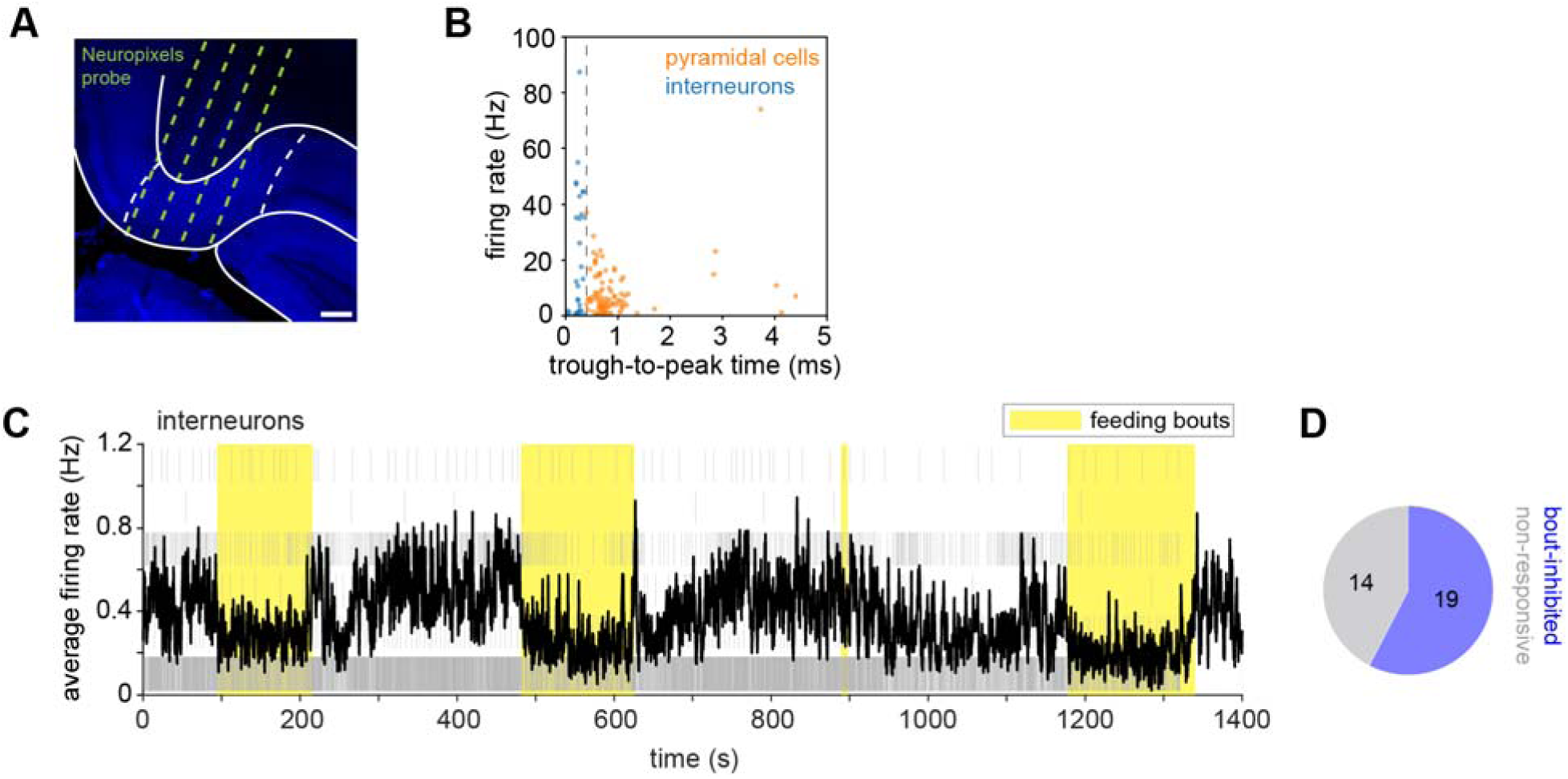
Additional characterization of dSub Neuropixels 2.0 recording. (A) Representative histology showing Neuropixels 2.0 probe implant. Scale bar is 200 μm. (B) Criteria of classification of pyramidal neurons and interneurons. Sorted units with trough-to-peak time (TTP) < 0.4 ms are assigned to interneurons, and units with TTP > 0.4 ms are assigned to pyramidal neurons. (C) Raster plot and populational average firing rate of all interneurons from a representative mouse. For readability given the long time window, only every 20th spike is displayed in the raster plot. (D) Proportion of bout-inhibited and non-responsive interneurons. Total of 33 neurons from 4 mice.

**Fig. S3.**
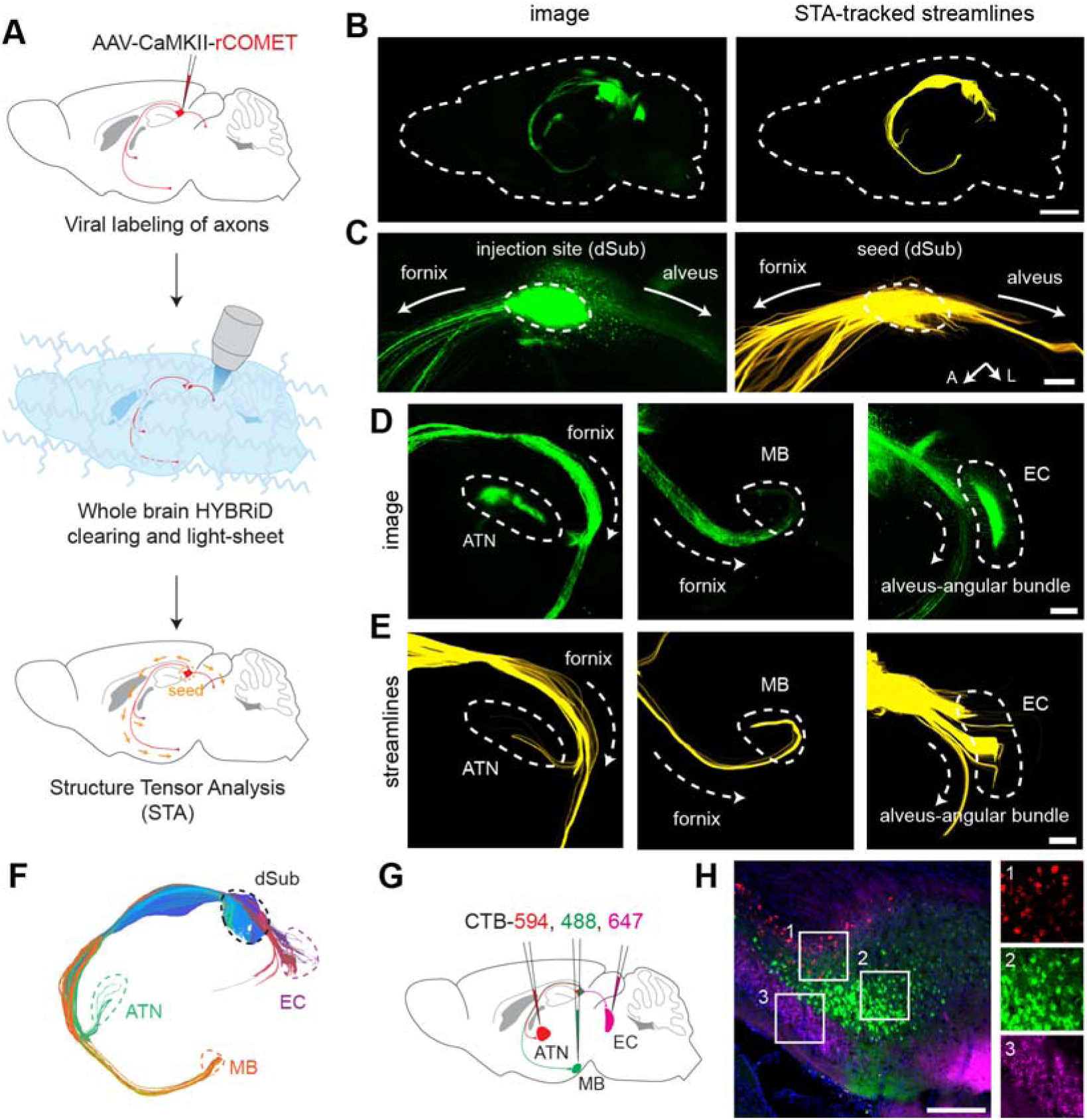
Mapping of extrahippocampal projections from dSub. (A) Schematic of virus injection, tissue clearing and light-sheet imaging, and structure tensor analysis. (B) Representative whole-brain image and reconstructed streamlines (direction of greatest tract diffusion). (C) Representative image and reconstructed streamlines around the dSub, showing two main streams of projections (fornix and alveus). (D-E) Representative images and reconstructed streamlines around ATN, MB and EC. (F) Reconstructed streamlines from dSub, colored by projection terminals. (G) Schematic of 3-color CTB retrograde tracing from ATN, MB and EC. (H) Representative images of dSub, showing distinct soma localizations of cells projecting to different targets. Scale bars are 1 mm for (B), and 200 μm for (C-E) and (H).

**Fig. S4.**
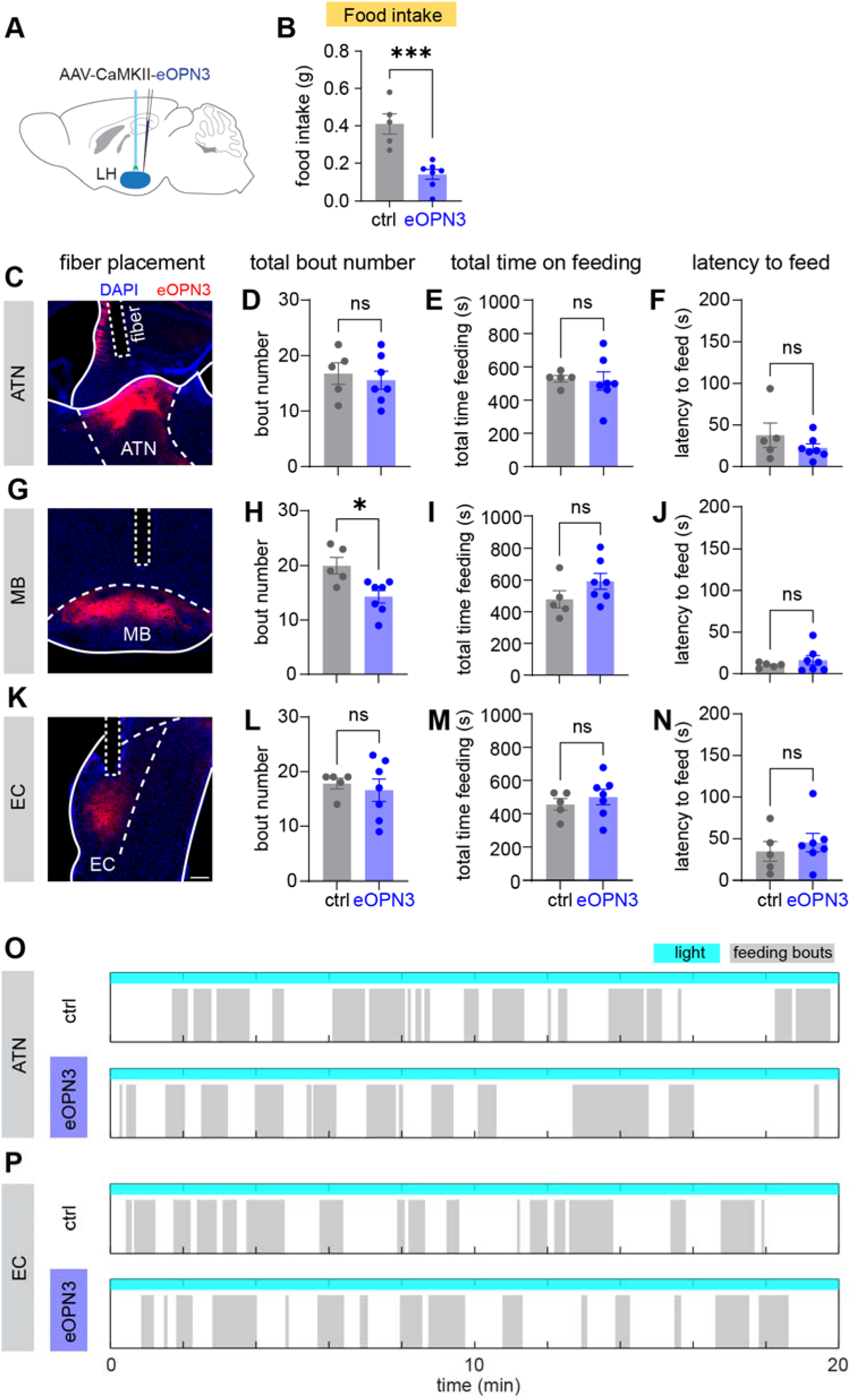
Additional characterization of feeding microstructure of dSub projection-specific photoinhibition. (A) Schematic of virus injection and fiber implantation for LH photoinhibition experiment. N = 5 mice for ctrl group and N = 7 mice for eOPN3 group. (B) Food intake analysis in the LH photoinhibition experiment. (C) Representative histology of ATN of dSub-ATN eOPN3 cohort, showing fiber placement and expression of eOPN3 in axons in ATN. (D) Total number of feeding bouts under dSub-ATN photoinhibition. (E) Total time spent in feeding bouts under dSub-ATN photoinhibition. (F) Latency to the first feeding bout under dSub-ATN photoinhibition. (G-J) Representative histology and quantifications for dSub-MB photoinhibition experiment. (K-N) Representative histology and quantifications for dSub-EC photoinhibition experiment. (O-P) Representative raster of feeding bouts of a ctrl mouse and an eOPN3 mouse of dSub-ATN and dSub-EC photoinhibition. All values are mean ± SEM. Statistics determined by Student’s t-test for mean duration and food intake analyses, and two-tailed permutation KS test for ECDF analyses. ∗p < 0.05; ∗∗∗p < 0.001. All scale bars are 200 μm.

**Fig. S5.**
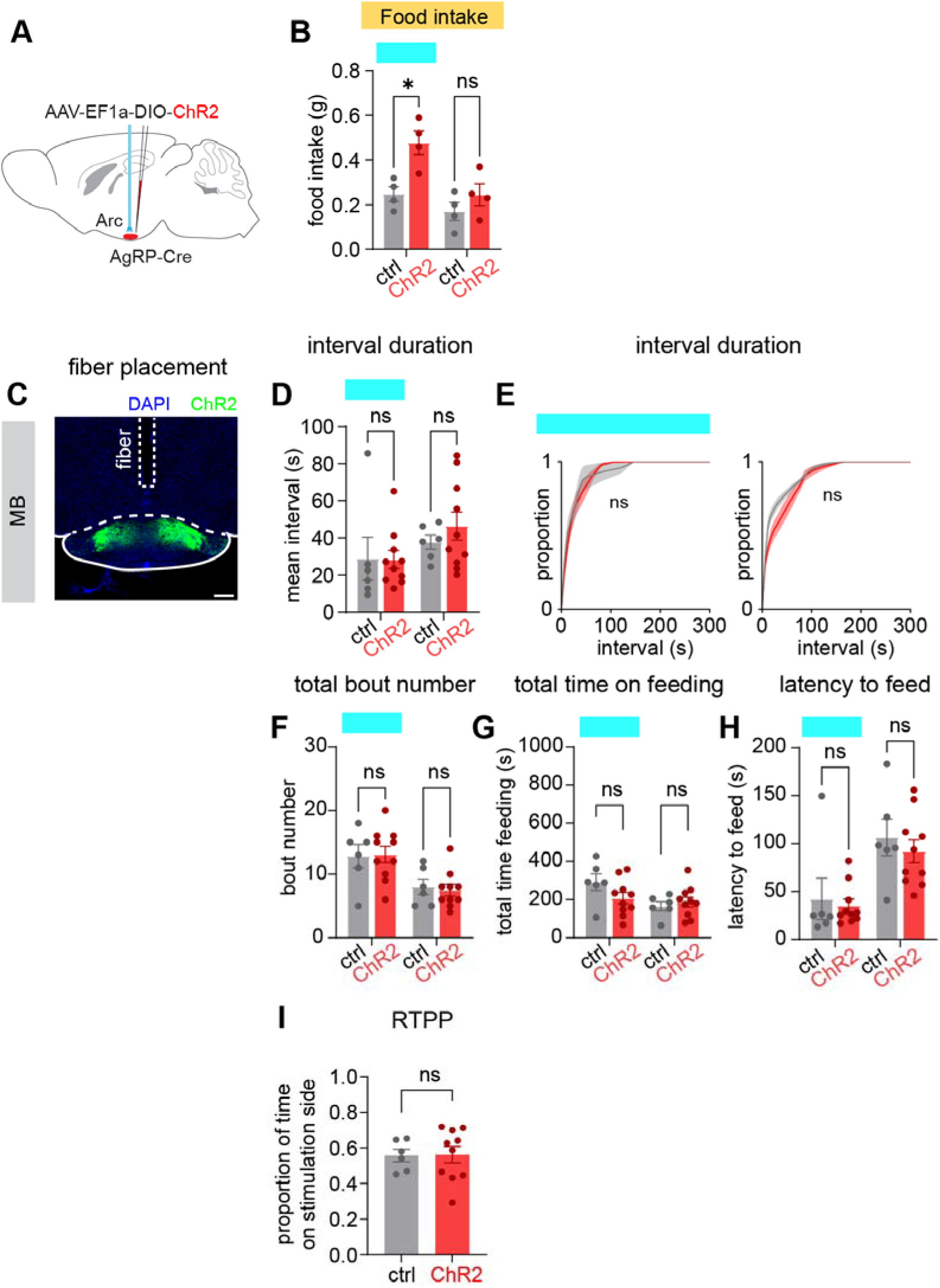
Additional characterization of feeding microstructure of dSub-MB photo activation. (A) Schematic of virus injection and fiber implantation for Arc AgRP photoactivation experiments. (B) Total food intake amount under Arc AgRP photoactivation. (C) Representative histology of MB from dSub-MB ChR2 cohort, showing fiber placement and expression of ChR2 in axons in MB. (D-E) Mean and ECDF of interval duration in light ON phase and light OFF phase of dSub-MB photoactivation. (F-H) Total feeding bout count, total time spent in feeding bouts, and latency to the first feeding bout under dSub-MB photoactivation. (I) Time spent on the stimulation side in a real-time place preference (RTPP) experiment. All values are mean ± SEM. Statistics determined by Student’s t-test for mean duration and food intake analyses, and two-tailed permutation KS test for ECDF analyses. ∗p < 0.05. Scale bar is 200 μm.

**Fig. S6.**
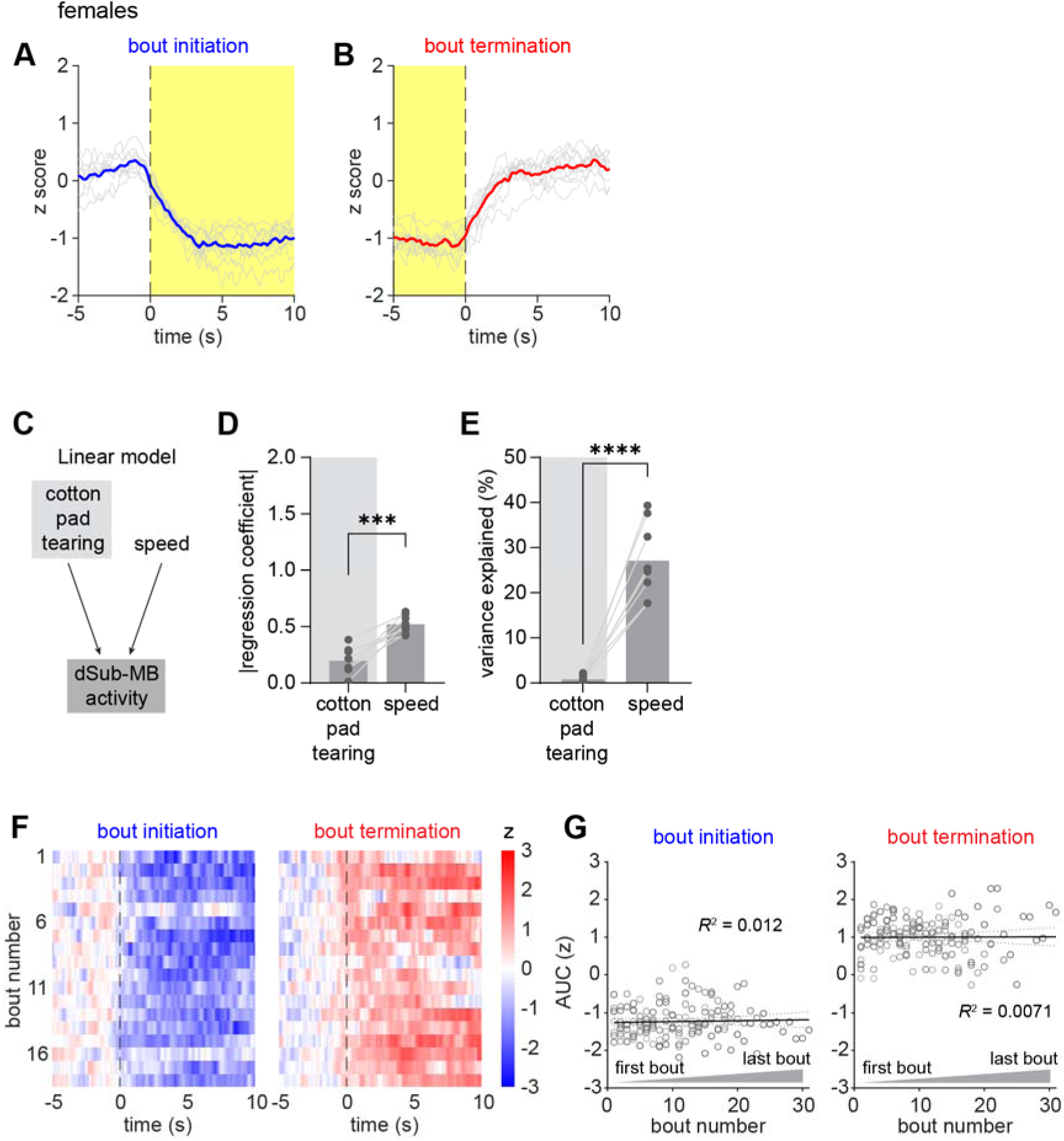
Calcium response of dSub-MB circuit during feeding and movement-related behaviors. (A-B) Averaged calcium signal in female mice triggered on initiation and termination of feeding bouts. N = 10 mice. (C) Schematic of the linear regression of dSub-MB calcium activity against bouts of cotton pad tearing and speed of locomotion. (D-E) Absolute value of regression coefficient and percentage of variance explained of dSub-MB activity by cotton pad tearing and speed. (F) Representative heatmaps showing dSub-MB activity triggered on initiation and termination of feeding for all bouts from a given mouse. (G) Correlation between AUC of neural activity during a feeding bout and the bout number within a session. Statistics determined by paired t tests in (D-E) and mixed-effect linear regression in (G).

**Fig. S7.**
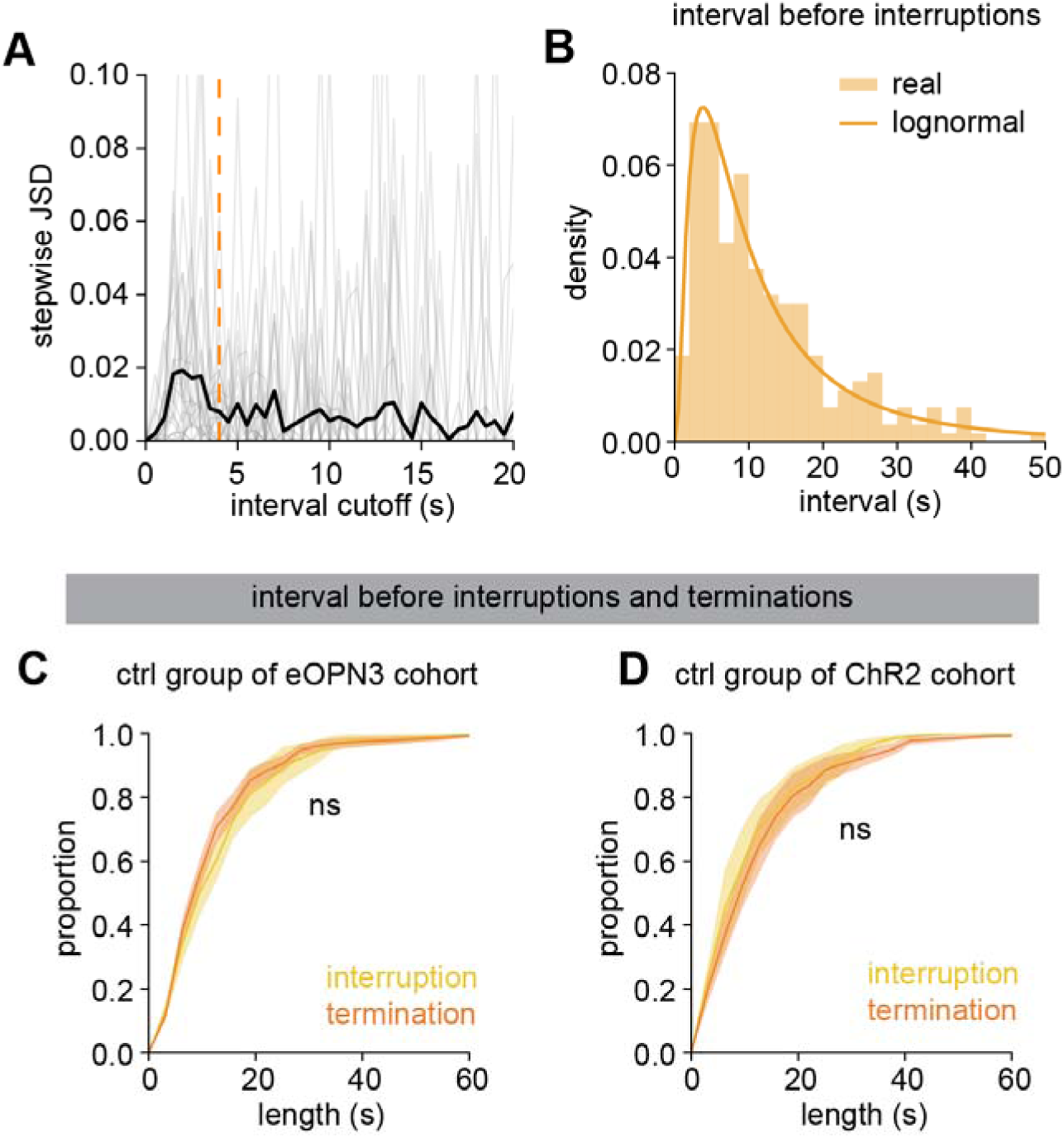
Additional characterizations of the interruptions. (A) Selection of a threshold to distinguish feeding bout interruptions from feeding bout terminations, and its effect on the distribution of feeding bout durations. Y axis shows difference in the empirical distribution of bout durations given a cutoff of t vs t + 0.5s, quantified by the Jensen–Shannon divergence (JSD). Orange dashed line shows the used cutoff at 4 s. Gray traces show individual mice, and black shows the average across all mice. N = 12 mice, combining all ctrl mice from the eOPN3 and ChR2 cohorts. (B) Histogram of the intervals between each intra-bout interruption or bout termination and the preceding intra-bout interruption (or bout start) in the experimental data (bars), and the fitted lognormal distribution used to generate interruption input pulses in the model. Data from the ctrl groups of the eOPN3 and the ChR2 cohorts. (C) ECDF of the interval between each intra-bout interruption or bout termination and the preceding intra-bout interruption (or bout start) for the ctrl group in the eOPN3 cohort. (D) ECDF of the interval between each intra-bout interruption or bout termination and the preceding intra-bout interruption (or bout start) for the ctrl group in the ChR2 cohort. Statistics determined by two-tailed permutation KS tests in (C-D).

**Fig. S8.**
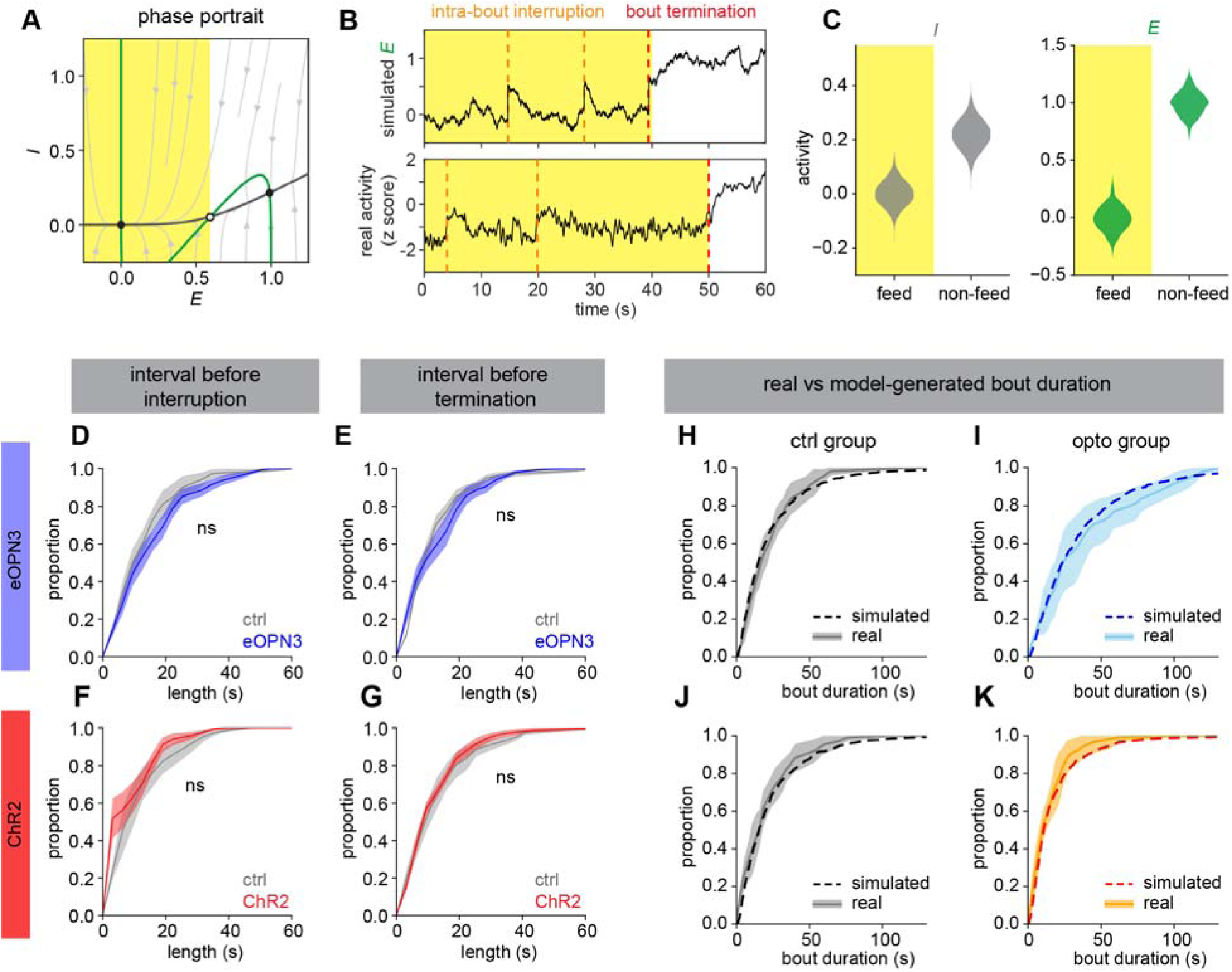
Additional characterizations of the model. (A) Phase portrait of the model. Nullclines of E and I populations are shown by the green and black lines, respectively, and flow field are shown by the light gray arrows. Solid dots are stable fixed points, hollow dots are unstable fixed points. We use the E population state at the unstable fixed point as the approximate threshold between states; sub-threshold E population values correspond to the feeding state, super-threshold to the non-feeding state. (B) Simulated activity of the E population (above) compared to fiber photometry recording of dSub-MB projection neurons from a representative mouse (below). (C) Activity distribution of the model I population and E population in the feeding and nonfeeding states. (D) ECDF of the interval between each intra-bout interruption and the preceding intra-bout interruption (or bout start) for the ctrl group in the dSub-MB photoinhibition cohort. (E) ECDF of intervals preceding a termination for the ctrl group and eOPN3 group in the dSub-MB photoinhibition cohort. (F-G) Same ECDFs for the ctrl group and ChR2 group in the dSub-MB photoactivation cohort. (H-I) ECDF of bout durations in real data and model-simulated data, for the eOPN3 cohort. (J-K) ECDF of bout duration from real data and model-simulated data, of the ChR2 cohort. Values are mean ± SEM for (D-G) and mean ± SD for (H-K). Statistics determined by two-tailed permutation KS tests for (D-G).

**Fig. S9.**
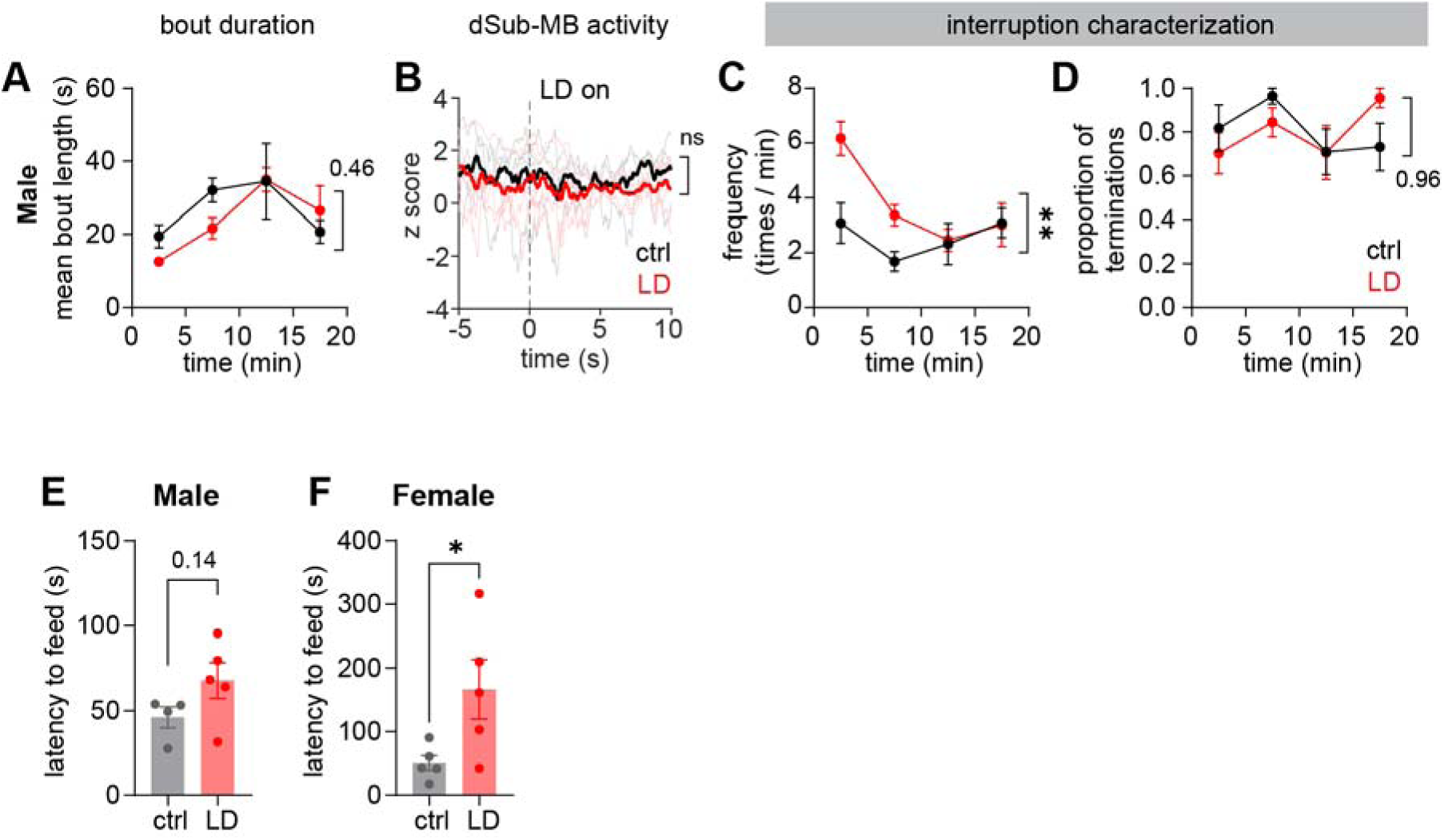
Additional characterizations of looming disc experiment. (A) Time course of the bout duration in male mice. N = 4 mice for ctrl group and N = 5 mice for LD group. (B) Calcium activity of dSub-MB in female mice triggered on the first appearance of the looming disk. (C-D) Time course of the total frequency of interruptions and terminations, and percentage of terminations, in male mice. (E-F) Latency to the first feeding bout in male and female mice, in control conditions vs in the presence of a looming disc (LD). All values are mean ± SEM. Statistics determined by two-way ANOVA (main effect of group) for (A-D), and Student’s t tests for (E-F). ∗p < 0.05, ∗∗p < 0.01. P values for non-significant comparisons are labeled.

